# Experimental and Bioinformatic upgrade for genome-wide mapping of nucleosomes in *Trypanosoma cruzi*

**DOI:** 10.1101/2021.07.02.450927

**Authors:** Paula Beati, Milena Massimino Stepñicka, Salomé Vilchez Larrea, Guillermo Daniel Alonso, Josefina Ocampo

## Abstract

In *Trypanosoma cruzi*, as in every eukaryotic cell, DNA is packaged into chromatin by octamers of histone proteins that constitute nucleosomes. Besides compacting DNA, nucleosomes control DNA dependent processes by modulating the access of DNA binding proteins to regulatory elements on the DNA; or by providing the platform for additional layers of regulation given by histone variants and histone post-translational modifications. In trypanosomes, protein coding genes are constitutively transcribed as polycistronic units. Therefore, gene expression is controlled mainly post transcriptionally. However, chromatin organization and the histone code influence transcription, cell cycle progression, replication and DNA repair. Hence, determining nucleosome position is of uppermost importance to understand the peculiarities of these processes in trypanosomes. Digestion of chromatin with micrococcal nuclease followed by deep sequencing has been widely applied for genome-wide mapping of nucleosomes in several organisms. Nonetheless, this parasite presents numerous singularities. On one hand, special growth conditions and cell manipulation are required. On the other hand, chromatin organization shows some uniqueness that demands a specially designed analytical approach. An additional entanglement is given by the nature of its genome harboring a large content of repetitive sequences and the poor quality of the genome assembly and annotation of many strains. Here, we adapted this broadly used method to the hybrid reference strain, CL Brener. Particularly, we developed an exhaustive and thorough computational workflow for data analysis, highlighting the relevance of using its whole genome as a reference instead of the commonly used Esmeraldo-like haplotype. Moreover, the performance of two aligners, Bowtie2 and HISAT2 was tested to find the most appropriate tool to map any genomic read to reference genomes bearing this complexity.

## 2 Introduction

In eukaryotes the nuclear DNA is packaged into chromatin, which is organized in repetitive units called nucleosomes. These basic units are composed of two of each of the canonical histones: H2A, H2B, H3, and H4, constituting a protein core around which a stretch of ~147 bp of DNA is wrapped (Luger et al., 1997).

*In vivo*, nucleosomes are regularly spaced by linker DNA. There seems to exist a correlation between DNA accessibility and the linker length, suggesting that nucleosome spacing is the primary determinant of gene regulation (Chereji et al., 2019). Additionally, a primary role of nucleosomes in controlling transcription was recently proposed (Kornberg and Lorch, 2020). Hence, studying nucleosome positioning is of paramount relevance to understand how genomic DNA is packed and how DNA-dependent processes are regulated.

In *T. cruzi*, chromatin is also organized in repetitive units (Astolfi et al., 1980) but shows some unusual characteristics compared to other organisms. Among these peculiarities, 30 nm fibers do not form *in vitro* (Hecker and Gander, 1985) and chromatin does not condense into chromosomes during mitosis (Hecker et al., 1994). This phenomenon is suspected to be due to the unique characteristic of its histone H1, which lacks the globular domains typically present in other eukaryotes (Toro and Galanti, 1988). Besides, the infective and non-infective parasite life forms display different levels of chromatin condensation during interphase (Hecker and Gander, 1985), differential susceptibility to DNaseI (Spadiliero et al., 2002) and some differences in the nucleosome landscape (Lima et al., 2021). Additionally, the histones of trypanosomes are the least conserved among all eukaryotic histones studied so far, differing from other organisms in size, sequence, and charge (Toro and Galanti, 1988; Thatcher and Gorovsky, 1994). Given all these distinctive properties and the potential implications for key cellular processes, a dedicated study of *T. cruzi* chromatin organization is required.

Several methods for genome-wide high-resolution chromatin profiling have been developed but not all of them are easy to adapt to every organism under study (Clark, 2010; Chereji and Clark, 2018). Despite having some limitations and caveats, the most widely used method, applied first in yeast, is the digestion of chromatin with micrococcal nuclease (MNase) followed by deep sequencing using paired-end technology (MNase-seq)(Cole et al., 2012). In the last few years, a huge number of nucleosome maps of numerous organisms including *T. brucei* and *Leishmania major* were generated (Lombraña et al., 2016; Maree et al., 2017; Wedel et al., 2017). More recently, a good experimental setup for epimastigotes and trypomastigotes of the *T. cruzi* CL Brener strain was published (Lima et al., 2021). However, given the hybrid nature of the CL Brener strain several analytical challenges remained unsolved.

Despite the large number of informatic tools currently available (Teif, 2016), a customized informatic workflow is required. All *T. cruzi* strains are diploid but there are two kinds: (1) Clonal strains, harboring equivalent homologous chromosomes; and (2) hybrid strains, carrying one copy of each chromosome from two substantially different parental strains. The latter adds some complexity to the data analysis. In this sense, numerous genomic studies are performed using the reference hybrid strain CL Brener, composed of the Esmeraldo-like and non Esmeraldo-like haplotypes. However, for data processing the community working in the field sometimes uses just one haplotype as reference genome for simplicity. In this work, a detailed protocol for nucleosome preparation and genome-wide mapping for *T. cruzi* epimastigotes is presented. Moreover, a thorough analysis was performed to find the most suitable steps to cope with the special challenges imposed by the hybrid nature of CL Brener, emphasizing the importance of using its whole genome as a reference. Furthermore, an itemized pipeline for analytical processing is provided and easy to adapt to any *T. cruzi* strain.

## 3 Materials and Method

### 3.1 General outline

The main objective of this method is to obtain fragments of ~147 bp of DNA, which are protected from MNase digestion by the presence of the nucleosome core. Exponentially growing *T. cruzi* epimastigotes are collected, permeabilized and chromatin is digested with MNase. After stopping the reaction, DNA is extracted, and the level of digestion is checked in an agarose gel. MNase-seq libraries are constructed as described (Ocampo et al., 2016, 2019). Due to the peculiarities of *T. cruzi* and the unprecedented analysis with this parasite, we follow the standard bioinformatic analysis described for yeast (Chereji et al., 2017; Beati and Chereji, 2020). Nevertheless, we tested alternative strategies at every step of the way to find the most suitable path (details below). Experimental and informatics workflows are shown in Figure 1A and B respectively.

**Figure 1.**
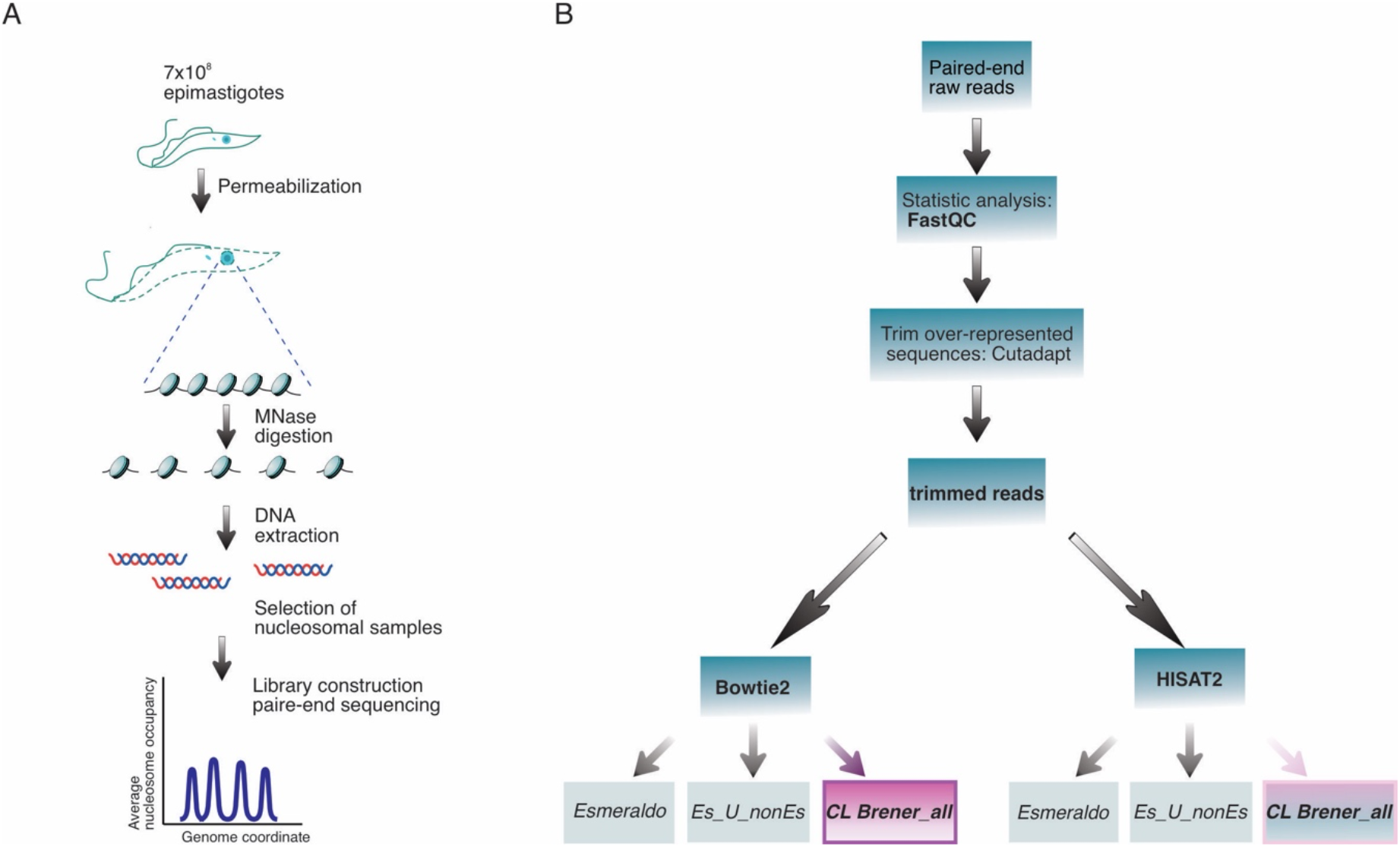
Experimental and informatic workflow. **(A)** Schematic representation of the experimental workflow. **(B)** Schematic workflow of the pipeline used to analyze the MNase-seq data. The use of the *CL Brener_all* as reference genome is suggested independently of the aligner of choice.

### 3.2 Epimastigote cultivation and harvest

CL Brener epimastigotes were grown at 28 °C in liver infusion tryptose (LIT) medium (5 g.l^-1^ liver infusion, 5 g.l^-1^ bacto-tryptose, 68 mM NaCl, 5.3 mM KCl, 22 mM Na_2_HPO_4_, 0.2% (w/v) glucose, and 0.002% (w/v) hemin) supplemented with 10% (v/v) fetal calf serum (FCS), 100 U/ml penicillin and 100 mg. l^-1^ streptomycin. Cell density was maintained between 1×10^6^ and 1×10^8^ cells.ml^-1^ by sub-culturing parasites every 6-7 days.

### 3.3 MNase digestion of *T. cruzi* epimastigote chromatin

Late exponentially growing epimastigotes of the CL Brener strain were collected. In each experiment, 7×10^8^ parasites were pelleted by centrifugation at 1700 xg for 5 minutes at room temperature (RT). Supernatant was discarded and the pellet was washed with 10 ml Incomplete Lysis Solution (1 mM L-glutamine, 250 mM sucrose, 2.5 mM CaCl_2_, 1 mM phenylmethylsulfonyl fluoride (PMSF), followed by centrifugation at 1700 xg for 5 minutes. After centrifugation, the supernatant was discarded. Then, cells were permeabilized as previously described with modifications (Leandro de Jesus et al., 2017). Briefly, cells were lysed with 800 μl Complete Lysis Solution (10 ml incomplete Lysis Solution, 0.01% Triton X 100, 1 mM PMSF). Then, samples were divided into 7 aliquots of 100 μl each into 1.5 ml microcentrifuge tubes, to test increasing amounts of MNase for a fixed period of time. The aliquots were centrifuged at 1700 xg for 5 minutes and the supernatant was discarded. The pellets were washed with 100 μl Incomplete Lysis Solution followed by centrifugation at 1700 xg for 5 minutes. To perform the MNase digestion, each pellet was resuspended in 400 μl of MNase Digestion Buffer (10 mM HEPES, 35 mM NaCl, 500 μM MgCl_2_, 500 μM CaCl_2_, 5 mM 2-mercaptoethanol and protease inhibitors). The buffer was prewarmed in a 25 °C water bath for 2 min before resuspending the samples. The digestion was started by the addition of increasing amounts of MNase (10U/μl) to each tube, for example: 0, 1, 3.5, 7, 10, 20, 40 μl. Then, the treated aliquots were mixed thoroughly without vortexing and incubated at 25 °C for 5 min. Reactions were stopped by adding 50 μl fresh MNase Stop Solution (180 mM EDTA, 7.2% SDS prepared in H_2_O) to each digestion, mixing, and leaving at room temperature for 5 min. After stopping the reaction, 50 μl 10% SDS (final concentration ~1.8% in ~500 μl) and 250μl of 3M KOAc (~1M final concentration in ~750μl) were added to each aliquot and mixed. Then, DNA was extracted by adding 750 μl chloroform, mixing and spinning in a microcentrifuge (5 min at top speed). DNA extraction was repeated once: supernatants were transferred to 1.5-ml tubes, 750 μl chloroform was added, mixed, and spun down in a microcentrifuge (5 min at top speed). Supernatants were transferred one more time to new 1.5 ml tubes and 0.7 vol. isopropanol was added to precipitate the DNA. The tubes were left at −20°C overnight. DNA was precipitated by spinning in a microcentrifuge (30 min at top speed). The supernatant was discarded, and the pellet was washed once with 0.6 ml 70% ethanol and spun again in a microcentrifuge (5 min at top speed). After centrifugation the supernatant was discarded. Each pellet was air-dried and dissolved in 50 μl of 10 mM Tris HCl pH 8, 2 mM EDTA with 100 μg/ml of RNase and incubated at 37 °C for 2 hours. **Note:** We recommend preparing MNase stock solution as follows: Dissolve MNase (Worthington) to 10 U/ml in 5 mM Na-phosphate pH 7.0, 0.025 mM CaCl_2_. Store in aliquots at −80 °C

### 3.4 Selecting the best mononucleosomal sample to proceed with

To determine the extent of digestion 10 μl of each MNase-digested sample was analyzed in a 2% agarose gel next to the PCR DNA Marker (New England Biolabs, Ipswich, Massachusetts, U.S.A.). An ideal sample should have more than 80% of the DNA represented in the mononucleosome band and a faint band in the di-nucleosome band as shown in Figure 2. In poorly digested samples a ladder of bands is observed, and only a small fraction of the total DNA is contained in the mononucleosome fraction. This mononucleosome band is longer than 150 bp due to poor trimming. Moreover, early digestion is biased to AT-rich DNA sequences since they are preferentially cut by MNase and the protected regions may include non-histone DNA binding complexes (Chereji et al., 2017). On the other hand, in over digested samples, faint bands appear underneath the mononucleosome band due to excessive trimming of the core particle. It is still possible to tell the position of the remaining nucleosomes, but some of them will be lost. Therefore, nucleosome occupancy maps will not be accurate. Hence, a compromise is required, keeping in mind that ~80% of the DNA should be contained in the mononucleosome band. Testing an aliquot of the selected sample in an Agilent Bioanalyzer 2100 expert with DNA 1000 (Agilent Scientific Instruments, Santa Clara, CA, US), which provides electropherograms and gel-like images, might help to make the correct decision, but is not indispensable. In any case, the precise length distribution of the chosen sample will be verified after pair-end sequencing as described below.

**Figure 2.**
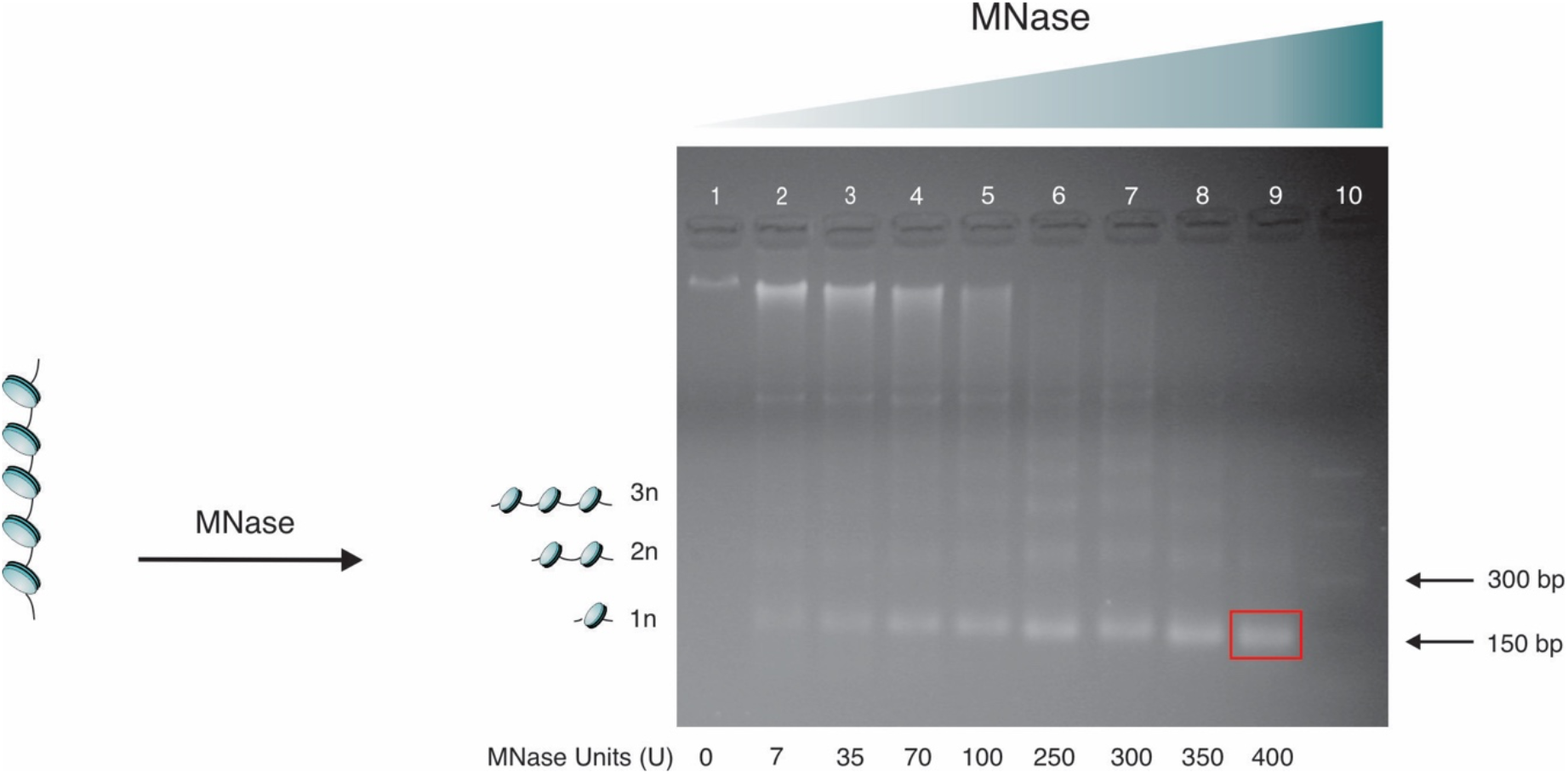
Titration of *T. cruzi* chromatin with MNase. Chromatin digested with increasing amounts of MNase analyzed in a 2% agarose gel (lanes 1-9). An example of the level of digestion of choice is highlighted in lane 9. PCR DNA marker (NEB) was loaded in lane 10.

### 3.5 Paired-end sequencing

After selecting the sample with the most appropriate level of digestion, paired-end libraries were prepared as described before (Ocampo et al., 2016, 2019).

In earlier work, samples were purified from a gel before library preparation and paired-end sequencing (Cole et al., 2012). However, purification is not necessary given that size sorting can be done *in silico* afterwards. Moreover, it can be useful to count on the whole data including additional DNA lengths (e.g.: di-nucleosome bands) present in the sample for further analysis (Ocampo et al., 2019).

Briefly, once the DNA samples with the appropriate level of digestion were selected, any nicks were repaired using the PrePCR kit (New England Biolabs, Ipswich, Massachusetts, US), purified using a Qiagen PCR column (Hilden, Germany) and eluted with 50 μl TE buffer (10 mM Tris-Cl pH8, 0.1 mM EDTA). Modification of the 5’ and 3’ ends of the nucleosomal DNA and adapter ligation were performed with the NEBNext Ultra DNA Library Prep Kit for Illumina and NEBNext Multiplex Oligos for Illumina (New England Biolabs, Ipswich, Massachusetts, US). Agencourt AMPure XP beads (Beckman-Coulter, Brea, California, US) were used to purify the DNA after adaptor ligation and PCR products before sequencing as described (Ocampo et al., 2016, 2019).

For sequencing reactions, either Illumina Next-seq 500 or Hi-seq 2000 were used to obtain pairs of 50 nucleotide (nt) reads.

### 3.6 Bioinformatic analysis

Basic bioinformatic analysis is straightforward when analyzing data from model organisms such as yeast, fly or mouse. However, *T. cruzi* presents so many peculiarities that extensive work is required. To find the most appropriate steps for *T. cruzi* analysis, several points of the standard workflow were tested. First, we analyzed whether it was necessary or not to trim over-represented sequences (ORS) from the fastq files. Second, the capacity to achieve the most accurate alignment was compared between the widely used aligner Bowtie2 (Langmead and Steven L Salzberg, 2013) and the newer tool HISAT2 (Kim et al., 2015). Third, alternative reference genomes were tested, including the commonly used Esmeraldo-like haplotype, a combination of both haplotypes (Esmeraldo-like + non Esmeraldo-like haplotypes) and a more comprehensive one including the non-designated regions (Esmeraldo-like + non Esmeraldo-like haplotypes + “extra”) (Figure 1B). Once the most reliable workflow was established, the pipeline to obtain length distribution histograms of DNA sequences, bigwig files, average plots and heatmaps were generated (see details below). Additionally, source code easy to adapt to any *T. cruzi* strain is available at (https://github.com/paulati/nucleosome).

#### 3.6.1 Data files and statistical analysis

Paired-end reads are normally provided in the fastq file format containing the raw reads. To do quality control checks on raw data, fastQC analysis was performed. During this step ORS were detected and trimmed out to avoid the generation of artefacts during the analysis. ORS in general are 50 nts long and correspond to adaptors or primers used for library construction. In the case of *T. cruzi*, due to the repetitive nature of its genome, some ORS occasionally contain runs of repetitive sequences. ORS were removed when present using the Cutadapt tool (Martin, 2011) producing trimmed fastq files, containing trimmed raw reads that from now on we will refer to as “*trimmed reads*”. Given that this step is regularly omitted when analyzing MNase-seq data from model organisms, to evaluate if it constitutes an improvement in the analysis of *T. cruzi*, we pursued the following steps in parallel using raw reads and trimmed reads.

#### 3.6.2 Sequence alignment and reference genome

To test which tool is more efficient to perform CL Brener alignments, Bowtie2 (Langmead and Steven L Salzberg, 2013) and the newer tool HISAT2 (Kim et al., 2015) were used to align either the raw or the trimmed 50 nt paired-end reads, to the corresponding reference genome. Bowtie2 is the most widely used aligner and is considered a multi-purpose tool; while HISAT2, was originally designed for RNA-seq analysis. HISAT2 bears a few optimizations including improving the accuracy of short-read alignment. Therefore, we pursued in parallel both strategies (Figure 1 B). To carry out this comparison, we set the same parameters values for both tools. In particular, both tools have a parameter called score-min, defined as a function of the read length, that is the minimum score needed for an alignment to be considered “valid”.

For Bowtie2, we used the end-to-end mode with default settings, commonly used for model organisms, where the score-min is specified by ‘L, −0.6, −0.6’. This, stablish a linear function f (x) = −0.6 + −0.6 * x, where x is the read length. In this case, x is 50 nts, which implies it accepts scores bigger than −30.6. In the case of Hisat2, the default settings are more restrictive than those for Bowtie2 with ‘L, 0, −0,2’. To make a fare comparison between both tools, we set for HISAT2 the exact same score-min we used for Bowtie2.

Due to the hybrid nature of CL Brener, its two haplotypes are annotated separately: Esmeraldo-like and non-Esmeraldo-like, each of them having 41 chromosomes, called S and P chromosomes respectively. An additional complication is that part of the genome is not assigned either to the Esmeraldo-like or to the non-Esmeraldo-like haplotypes and is annotated separately. In previous high throughput studies using CL Brener, the output data were aligned only to the Esmeraldo-like haplotype genome for simplicity (Lima et al., 2021). To test the noise introduced by this simplification, two additional reference genomes were generated for comparison. One, combining Esmeraldo-like and non-Esmeraldo-like haplotypes designated *Es_U_nonEs*; and the second one, combining Esmeraldo-like, non-Esmeraldo-like and the “extra” regions, designated *Cl Brener_all*. Versions TriTryp46 of the respective genomes were used as a reference for informatic analysis in every case.

#### 3.6.3 Binary file generation

Bigwig files containing information of nucleosome occupancy and nucleosome position were generated. Occupancy (coverage) maps were built by counting the number of times that a base pair was occupied by a nucleosome, while position maps counted the number of times each particular nucleosome sequence was present in the data set. The midpoint of the nucleosome, called the dyad, was represented as previously described (Clark, 2010; Cole et al., 2012; Chereji and Clark, 2018). To observe more clearly the nucleosomes in the occupancy maps and to be more accurate in the estimation of the position maps, the analysis was restricted to those fragments that belong to the nucleosome size range (120-180 bp) and is normalized by summing all the nucleosome sequences covering a nucleotide and dividing that number by the average number of nucleosome sequences per base pair across the genome.

#### 3.6.4 Prediction of the trans-splicing acceptor site

The 5’untranslated regions (5’UTR) were predicted with UTRme (Radío et al., 2018). Given the lack of transcriptomic data for CL Brener epimastigotes, the predictions were based on available RNA-seq data for the Y strain (Li et al., 2016) using TriTryp46 Esmeraldo-like genome as a reference. As an approximation of the trans-splicing acceptor site (TAS), the 5’ end of the 5’UTR region was used.

#### 3.6.5 Average plots and 2D occupancy plots

Average representation of the data and 2D Occupancy plots were performed as previously described (Chereji et al., 2017; Beati and Chereji, 2020) with some modifications. Average nucleosome determinations were performed from the alignments generated in all the alternative paths tested above (Figure 1B) and represented relative to the predicted TAS. Briefly, even when the total number of reads was previously aligned to the *Es_U_nonEs* or the *CL Brener_all* genomes, for this analysis only the average nucleosome occupancy coming from the reads that aligned to the Esmeraldo-like haplotype were represented relative to the predicted TAS. For 2D Occupancy plots, the same data was represented relative to the TAS in the x axis, while the size of the analyzed DNA fragments was represented in the y axis.

## 4 Results

### 4.1 Nucleosome preparation and deep sequencing

We established a straightforward protocol for nucleosome preparation from epimastigotes of any *T. cruzi* strain as described above. We achieved good quality nucleosome preparation for replicated experiments of CL Brener strains. An agarose gel for a representative experiment and the number of total paired-reads counted after aligning the replicated data to the reference genome is shown in Figure 3A and B respectively. The agarose gel for the replicate experiment is shown in Supplemental Figure 1.

**Figure 3.**
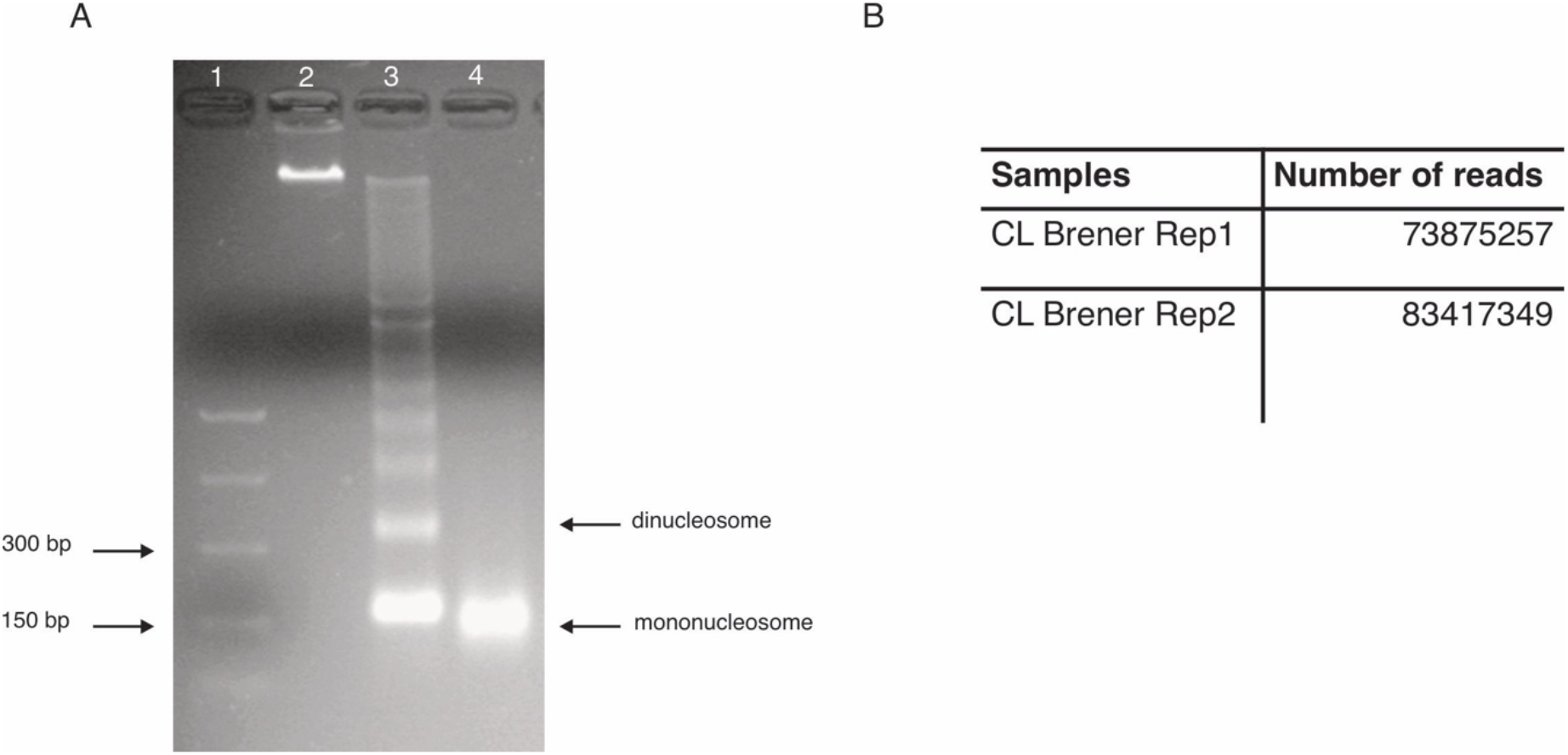
Mononucleosome sample and total number of paired-reads. **(A)** Chromatin digestions from the CL Brener replicate 1 experiment analyzed in a 2% agarose gel. The sample loaded in lane 4 was used for the experiment. PCR DNA (NEB) marker was loaded in lane 1. **(B)** Total number of paired-reads obtained in each replicate experiment.

### 4.2 Length distribution histograms

Once we aligned the paired-reads to the reference genome, we checked the level of digestion achieved for the sequenced samples by plotting a length distribution histogram, as previously described (Cole et al., 2012). Both samples from the two replicated experiments present a peak around the nucleosome size ~147-148 bp showing we obtained the right level of digestion (Figure 4). It is worth noting that to compare different samples, a good criterion is to use samples with similar length distribution histograms, indicating similar levels of digestion.

**Figure 4.**
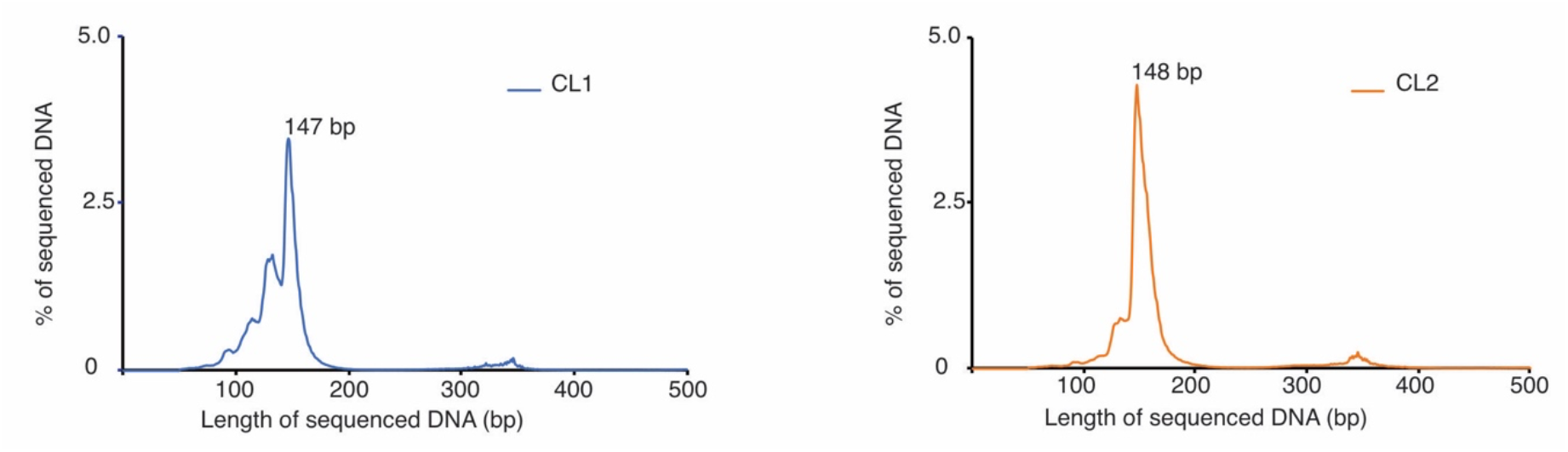
Length distribution of sequenced DNA. Length histogram for all nucleosomal DNA sequenced for replicate 1 (left panel) and replicate 2 (right panel) respectively.

### 4.3 Selection of the alignment tool and choice of the best reference genome for the CL Brener strain

To find the most suitable pipeline to analyze CL Brener data, we tested the alignment of raw and trimmed reads either to *Esmeraldo, Es_U_nonEs* or *CL Brener_all* genomes using Bowtie2 and HISAT2 (Figure1 B). We compared the output statistics of the alternative alignments for both replicated experiments, which are summarized in Table 1 and Supplemental Table1 respectively.

**Table 1.** Statistics of alignments. The percentage of alignments for the different paths tested with replicate 1 is summarized. Values obtained when using trimmed reads are coloured in green scale; values obtained when using raw reads are coloured in yellow scale. The higher the value the darker the colour.

|  |  |  |  |  |  |  |  |
| --- | --- | --- | --- | --- | --- | --- | --- |
| *CL Brener* | Bowtie2 | trimmed reads | genome | 0 times | 1 time | > 1 time | overall |
|  |  |  | *Esmeraldo* | 46.63 | 22.93 | 30.44 | 53.37 |
|  |  |  | *EsUnonES* | 36.42 | 10.66 | 53.1 | 63.76 |
|  |  |  | *CL Brener_all* | 28.89 | 9.05 | 62.06 | 71.11 |
|  |  | raw reads | genome | 0 times | 1 time | > 1 time | overall |
|  |  |  | *Esmeraldo* | 47.19 | 22.68 | 30.13 | 52.81 |
|  |  |  | *EsUnonES* | 36.67 | 10.36 | 52.96 | 63.33 |
|  |  |  | *CL Brener_all* | 29.58 | 8.9 | 61.52 | 70.42 |
|  | Hisat2 | trimmed reads | genome | 0 times | 1 time | > 1 time | overall |
|  |  |  | *Esmeraldo* | 47.01 | 41.65 | 11.34 | 52.99 |
|  |  |  | *EsUnonES* | 38.07 | 45.65 | 16.28 | 61.93 |
|  |  |  | *CL Brener_all* | 36.42 | 40.79 | 22.8 | 63.58 |
|  |  | raw reads | genome | 0 times | 1 time | > 1 time | overall |
|  |  |  | *Esmeraldo* | 47.58 | 41.2 | 11.21 | 52.42 |
|  |  |  | *EsUnonES* | 38.74 | 45.16 | 16.1 | 61.26 |
|  |  |  | *CL Brener_all* | 37.1 | 40.35 | 22.55 | 62.9 |

The first challenge was to test whether trimming the ORS was necessary, and we observed that in general it had a minimum effect for both aligners. When using Bowtie2, trimming raw reads only led to a tiny improvement in the overall alignment in less than 1% with any genome used (from 70.42% to 71.11% when using *CL Brener_all* genome; from 63.33% to 63.76 % when using *Es_U_nonEs* genome; or from 52.81% to 53.37% when using *Esmeraldo* genome), by slightly increasing the number of reads that align to the genome only one time, or more than once, accompanied by a small reduction in the number of reads that align zero times, consistent with the fact that most of the trimmed sequences correspond to adaptors or primer sequences. A similar trend is observed when using HISAT2 (Table1 and Supplemental table 1).

The second step was to evaluate the most suitable genome to use as a reference. For Bowtie2, the use of the *CL Brener_all* genome as a reference improved the overall alignment by ~7% with respect to the *Es_U_nonEs* genome (from 63.76% to 71.11%) and by ~18% with respect to the *Esmeraldo* genome (from 53.37% to 71.11%) when using trimmed reads at the expense of an increment in the number of reads that align more than once. This observation is associated with a significant reduction in the number of reads that align zero times, suggesting that the fraction of reads aligned only when using *CL Brener_all* belongs to regions of the genome that are not assigned to any of the haplotypes, probably due to their repetitive nature. While for HISAT2, using the *CL Brener_all* as the reference genome improves by only ~1.65% the overall alignment with respect to using *Es_U_nonEs* (from 61.93% to 63.58%), the improvement relative to using the *Esmeraldo* genome was ~10% (from 52.99% to 63.58%) based on reads that align more than once.

Third, comparing Bowtie2 with HISAT2, the overall alignment is better when using Bowtie2 by ~7.5 % when using the *CL Brener_all* genome (from 63.58% to 71.11%), by ~2 % when using the *Es_U_nonEs* genome (from 61.49% to 63.76%), but it is insignificant, when using the *Esmeraldo* genome (from 52.99% to 53.37%). Although the genome choice seems to be unimportant for HISAT2, when comparing *CL Brener_all* and *Es_U_nonEs*, using the *CL Brener_all* genome avoids artifactual alignments (discussed below). The biggest improvement of using HISAT2 over Bowtie2 is the increment in the number of reads that align only once by ~32% (from 9.05% to 40.79%) when aligning to *CL Brener_all* genome, by ~35% when aligning to *Es_U_nonEs* genome (from 10.66% to 45.65%), and by ^~^19% when using *Esmeraldo* (from 22.93% to 41.65%), consistent with HISAT2 being more efficient to detect uniquely aligned reads with any reference genome tested.

To summarize, trimming has a minimal effect in any path followed. However, considering that the removed reads are spurious, it is a good advice to cut them out to avoid any artifactual noise in later analysis. Overall, using Bowtie2 with *CL Brener_all* genome is a good choice and allows a better overall alignment than HISAT2. However, HISAT2 outstrips Bowtie2 by being more efficient to solve the alignment of reads at multiple places in the genome, particularly relevant when working with a hybrid strain (Table1 and Supplemental table 1). Finally, using *CL Brener_all* as a reference genome instead of *Es_U_nonEs*, or just *Esmeraldo* constitutes the critical point of the workflow (discussed in detail below).

### 4.4 Using *CL Brener_all* genome as a reference prevents spurious alignments

To make a visual inspection of how nucleosomes are organized in a particular region or loci of interest, we generated bigwig files containing either nucleosome occupancy maps or nucleosome position maps and inspected them in the Integrative Genome Viewer (IGV). This visualization facilitates the comparison between samples and the examination of nucleosome position or nucleosome occupancy maps against the chromosome coordinates. In Supplemental figure 2, we show nucleosome occupancy and nucleosome dyads for one representative region of CL Brener replicated experiments.

Additionally, we performed a comparison of the nucleosome occupancy generated from the different alignments described above using IGV. When we used either *Esmeraldo* or *Es_U_nonEs* as reference genomes with any aligner tested, we observed a few regions of extremely high nucleosome density (Figure 5A and Supplemental Figures 3A, 4A and 4B). These unusually high signals, that in general correspond to the most distal regions of the chromosomes, were smoothed out almost completely when using *CL Brener_all* as reference genome. Both aligners seem to work very well with the *CL Brener_all genome*; but when using HISAT2, we observed a couple of regions higher nucleosome density (Figure 5B and Supplemental Figures 3B and 4C). One of these unusually high signals was detected at the Mucin-associated surface protein (MASP) locus, a multi copy gene family. Finally, trimming had a minimum effect as observed in Figure 5B and Supplemental figure Figure 4C).

**Figure 5.**
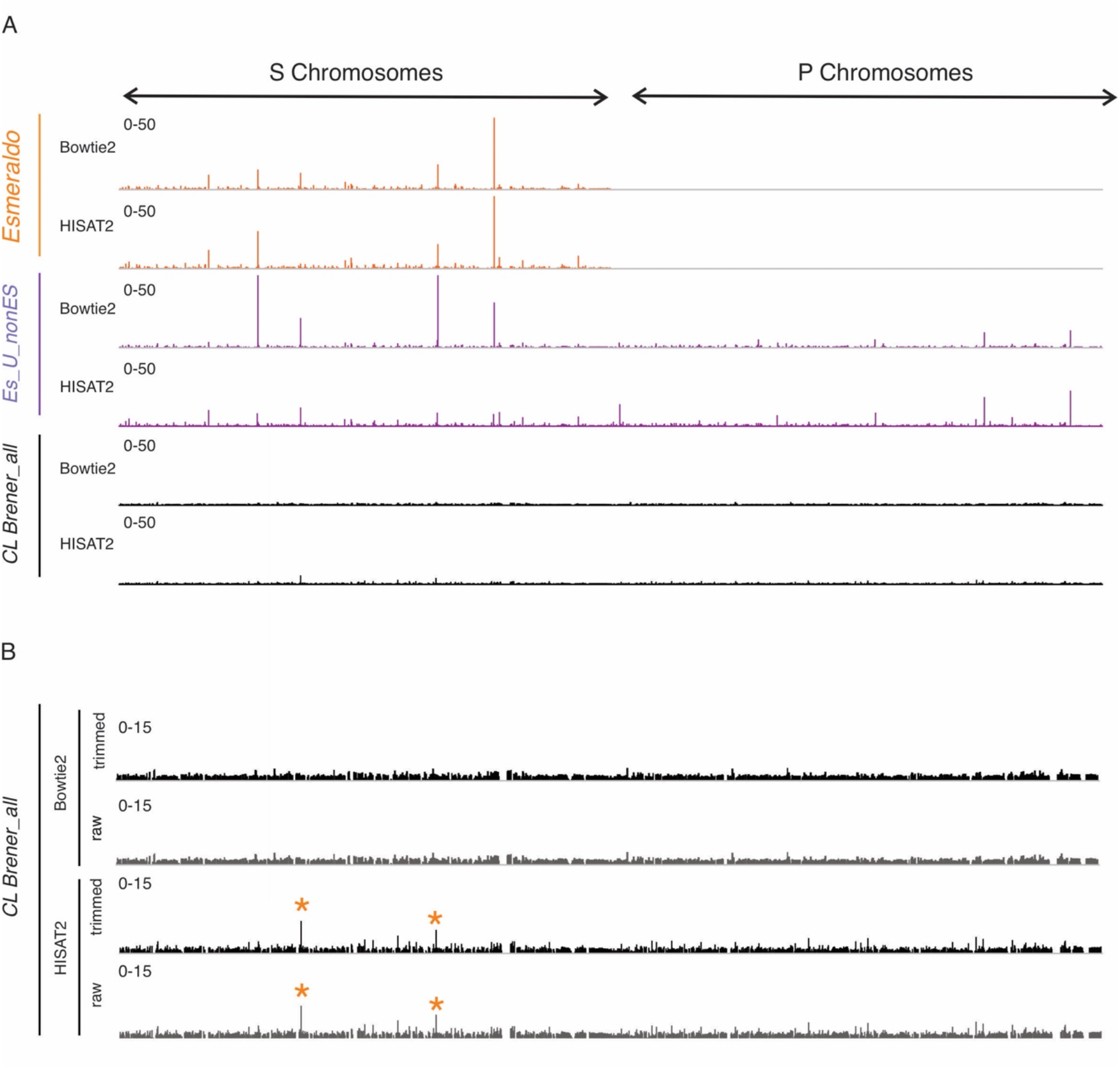
Using the CL *Brener_all* genome prevents spurious alignments. **(A)** IGV image for normalized nucleosome occupancy maps for the whole genome generated with trimmed reads from one representative data set (replicate 1) aligned to the *Esmeraldo* (orange), *Es_U_nonEs* (purple) and *CL Brener_all* (black) genomes respectively either with Bowtie2 or HISAT2. **(B)** Magnified IGV image for the nucleosome occupancy maps generated from trimmed and raw reads aligned to the *CL Brener_all* genome. Orange (*) indicates the main artefacts introduced when using HISAT2.

### 4.5 Average chromatin organization of the *T. cruzi* genome

For genomic analysis, manual viewer inspection of the data sets is not enough, it should be complemented with average analysis of the whole genome which is normally plotted relative to a reference point. For model organisms such as yeast, the average nucleosome occupancy or the average dyad occupancy for every gene in the genome is usually represented relative to the transcription start sites (TSS) or +1 nucleosome. This representation results in a prototypical chromatin organization with peaks at nucleosome midpoints and depressions at linker DNA. The peaks are regularly spaced and phased relative to the TSS and preceded by a nucleosome depleted region upstream of the +1 nucleosome (Yuan, 2005; Lee et al., 2007; Lantermann et al., 2010). This chromatin organization is broadly conserved among different organisms (Mavrich et al., 2008; Li et al., 2011; Valouev et al., 2011). However, the scenario is completely different for trypanosomes.

In Trypanosomatides, this genomic analysis is extremely challenging. On one hand, most of the genes are transcribed into polycistronic transcription units (PTUs) with no clear TSS. Each PTU matures into monocistronic mRNA by the addition of a 39 bp sequence denominated splice leader at the 5’UTR region and a 3’UTR polyadenylation in a process known as trans-splicing (Günzl, 2010).

In *L. major, T. brucei* and recently in *T. cruzi* poor chromatin organization was observed consistent with the fact that the genome is being constantly transcribed. In these organisms, nucleosomes are organized around the TAS, showing a small trough followed by a well-positioned nucleosome. However, in *T. brucei* internal TAS in the PTU differs from the first one by showing a well-positioned nucleosome with no nucleosome depleted region upstream (Lombraña et al., 2016; Maree et al., 2017; Lima et al., 2021). In this work, we improved the prediction of the TAS by using the UTRme predictor (Radío et al., 2018). as described above. We performed the TAS prediction only for the Esmeraldo-like haplotype. Then, we generated average nucleosome occupancy and 2D occupancy plots representing only the S chromosomes but using the bam files generated from all the alternative alignments described above (Figure 6 and Supplemental figures 5 and 6). As previously described, we observed a mild depression at the reference point in both the average and the 2D Occupancy plots. Moreover, the heatmaps allowed us to confirm that the size of the represented DNA is in the mononucleosome range. We obtained noiseless representations when using *CL Brener_all* genome either with Bowtie2 or HISAT2, and *Es_U_nonEs* genome with Bowtie2. However, when aligning to *Es_U_nonEs* with HISAT2 or to the *Esmeraldo* genome with either tool, some artefacts appeared, as denoted by the ladders of bands of high or low molecular weight in the heatmaps. These observations highlight the importance of using the *CL Brener_all* genome as a reference for cleaner results even when the analysis will be focused on only one haplotype.

**Figure 6.**
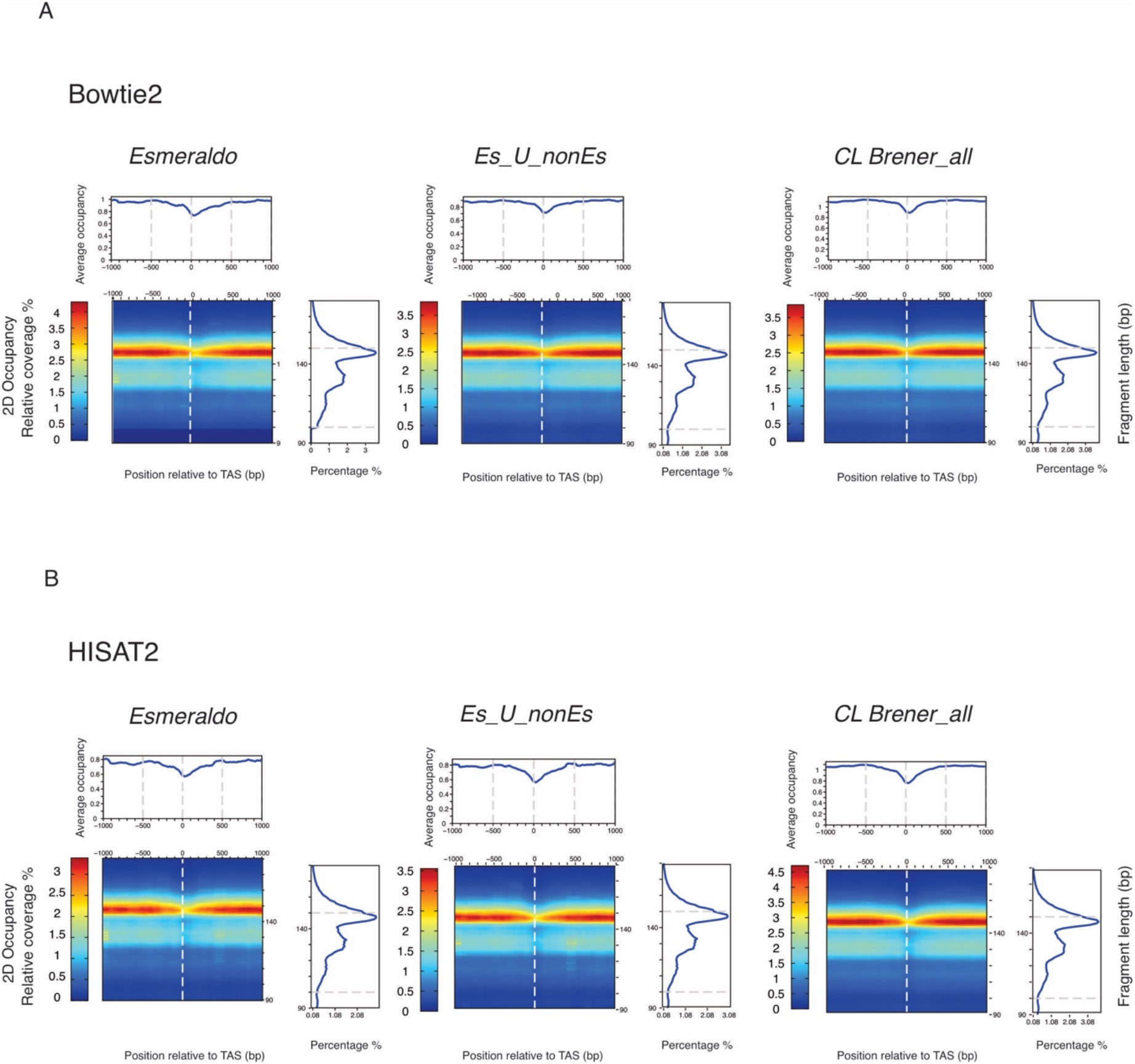
Using *CL Brener_all* genome prevents artifactual signals. Average nucleosome occupancy and 2D occupancy plots performed from trimmed reads for one representative data set (replicate 1) process with either **(A)** Bowtie2 or **(B)** HISAT2. Red: High nucleosome density; blue: low nucleosome density.

## 5 Discussion

In this work, we described a high-resolution method for genome-wide mapping of nucleosomes in *T. cruzi* epimastigotes providing an upgraded step by step workflow for the experimental approach. Moreover, we performed a categorical informatic analysis (Figure 1B). We have provided the experimental details and an informatic pipeline which are easily adaptable to any *T. cruzi* strain.

A good MNase-seq protocol for *T. cruzi* was previously published (Lima et al., 2021). Here, we upgraded it by providing cautious experimental details and careful considerations to perform the informatic analysis. In this regard, we first tested cell permeabilization as previously described for *T. brucei* using digitonin (Maree et al., 2017), but we encountered a huge variability in chromatin recovery among replicates. Hence, we switched to the above protocol. It is very important to avoid the use of a vortex at any step of the procedure. Every solution must be prepared in advance, except when using PMSF, 2-mercaptoethanol or protease inhibitors that should be added fresh to the solutions. The range of MNase to be tested is only a suggestion. It is recommended to perform a titration in every replicate experiment, as described in Figure 2, since the level of digestion achieved is somewhat unpredictable and depends on many variables such as permeabilization efficiency, reaction conditions and the experience of the operator.

To properly assign the location of nucleosomes in the genome, it is extremely important to use paired-end technology for DNA sequencing. The total number of paired-reads required to produce high resolution nucleosome occupancy maps depends on the size of the genome under study. For *T. cruzi*, it will depend on whether the strain is a clone or a hybrid. In the case of a hybrid strain like CL-Brener, the genome is estimated to be ~89.84 Mb and 41 pairs of chromosomes are designated (El-Sayed et al., 2005). Therefore, reads need to be split up into the two sets of chromosomes that comprise the hybrid. Hence, for global analysis, 10 or 15 million reads might be enough for clones and hybrid strains respectively, but for a more sophisticated approach, such as heatmap construction or sequence level analysis, a larger number of reads (~75 million) might be required, especially when working with hybrid strains. In this work, we obtained a remarkably good number of reads for each replicate experiment (Figure 3B). Although it could be tempting, replicate data should never be combined for analysis because comparison of biological replicates is essential to assess reproducibility.

Ideally, the strain of choice for high throughput studies should count on a well assembled and annotated genome. The first *T. cruzi* genome to be sequenced was the CL Brener strain (El-Sayed et al., 2005). Afterwards, the genomes of several other *T. cruzi* strains have been sequenced, assembled, and collected in the TriTryp database (https://tritrypdb.org/tritrypdb/). Unfortunately, most of them have been sequenced with short sequence read technologies, making accurate annotation almost impossible considering the high content of repetitive sequences of the *T. cruzi* genome. With the advent of long read technologies, such as Nanopore or PacBio, more reliable assemblies have been possible (Berná et al., 2018; Callejas-Hernández et al., 2018; Reis-Cunha and Bartholomeu, 2019). Unfortunately, at the onset of this work, no genome sequence for any *T. cruzi* strain was of outstanding quality. Hence, we chose CL Brener for being the reference strain and widely used by many researchers. Despite the of CL Brener genome being poorly assembled and annotated, we developed a careful workflow taking into account these caveats for cautious conclusions.

Accordingly, the data should be aligned to the best genome version of the strain in which the experiment was performed. Here, we showed that using only the Esmeraldo-like haplotype or even combined with the non Esmeraldo-like haplotype as reference genome, constitutes a blunder, introducing noise to the analytical process (Figure 5 and 6, and Supplemental figures 3, 4, 5 and 6). For in depth examination, it is a good decision to focus on the Esmeraldo-like haplotype since many studies and predictions from different labs have been based on this portion of the genome and it is useful for comparisons. However, the genomic data should be first aligned to the whole genome, *CL Brener_all*, as described in Figure 6 and Supplemental Figures 5 and 6.

Given the complex characteristics of the hybrid genome, the performance of the widely used aligner Bowtie2 was compared with HISAT2. Deciding which are the best parameters to use, represents a conundrum due to high content of repetitive sequences present in *T. cruzi* genome and the limited information about the exact percentage of these regions. From the statistics of alignments summarized in Table 1 and Supplemental Table 1) we observed that the overall alignment was somewhat higher when using Bowtie2, while HISAT2 was discarding some repetitive reads being more restrictive for ambiguous alignments. Particularly, HISAT2 was more efficient at resolving replicated reads providing a bigger proportion of uniquely aligned reads with any reference genome tested. Unfortunately, due to the uncertainty about the exact content of repetitive regions present in the CL Brener genome, it is almost impossible to ensure which aligner is achieving a more precise result. However, from the average analysis and heatmaps presented in this work (Figure 6 and Supplemental Figures 5 and 6), we concluded that both tools provide similar results when performing broad analysis. Additionally, careful examination on IGV shows that the alignment with HISAT2 results in a few regions show particularly high nucleosome density, even when choosing *Cl Brener _all* genome. These unusually high nucleosome density appears only in a few regions, in general coincident with multi copy gene families such as MASP, and most likely correspond to repetitive regions that were collapsed during genome annotation. Therefore, using either of these aligners with the *CL Brener_all* genome is a good choice, but their particular limitations might be taken into account for careful conclusions in each case. For global analysis Bowtie2 is the tool of choice for providing the biggest overall alignment, but when the analysis needs to be focus on uniquely aligned reads or single copy genes families HISAT2 could be an overcoming alternative.

## 6. Conclusion

In the last few years, the relevance of chromatin studies in trypanosomes has become more prominent. Here, we provide an updated experimental workflow and a robust informatic analysis. Our updated method not only gives a step by step experimental and analytical workflow for MNase-seq but is relevant to any high throughput study performed with the CL Brener strain.

It has been described that at least 50% of the CL Brener genome contains repetitive sequences and the two haplotypes differ by more than 5%. Additional work is still needed to improve the accuracy of genome assembly and annotations for CL Brener and other *T. cruzi* strains to get more certainty in the analysis of hithroughput studies.

## Supporting information

Suppemental_Material

## List of abbreviations

MNase: Micrococcal nuclease
ORS: overrepresented sequences
5’UTR5: 5’end of the 5’untraslated region
PTU: polytransciption unit
FCS: fetal calf serum
PMSF: phenylmethylsulfonyl fluoride
HEPES: 4-(2-hydroxyethyl)-1-piperazineethanesulfonic acid
EDTA: Ethilendiamine tetracetic acid
2ME: beta-mercaptoethanol
nts: nucleotides
bp: base pairs

## DATA AVAILABILITY STATEMENT

The datasets generated during the current study are available in the NCBI Gene Expression Omnibus (GEO) repository [GSE176341].

## Author contributions

JO and GDA conceived the study, designed the experiment and mentored the project.

MMS performed the experiments and contributed to figure design. PB performed data analysis and developed the adapted bioinformatic tools. JO collaborated in data analysis, made the figures and wrote the manuscript. SVL contributed to experimental setups. JO, GDA and PB carried out data interpretation.

## Acknowledgments

We are grateful to Dr. David Clark for valuable discussions and helpful comments on the manuscript. We thank Dr. Razvan Chereji for his informatic support at early stages of the project and helpful discussions during its development. We Thank Dr. Luis Diambra for suggestions on analytical tools and useful discussions. We thank the NHLBI Core Facility (Yan Luo, Poching Liu and Jun Zhu) for paired-end sequencing.

