## Supplementary material for "Experimental and Bioinformatic upgrade for genome-wide mapping of nucleosomes in *Trypanosoma cruzi*": Suppemental_Material

**Supplemental Figures and Table**

**Supplemental Figure 1 (associated to figure 3)**

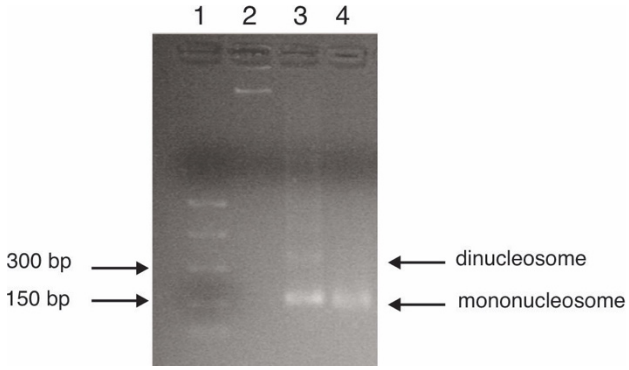


**Supplemental Figure 1.** Mononucleosome sample for replicate experiment. Chromatin digestions from epimastigotes of the CL Brener strain replicate 2 analyzed in a 2% agarose gel. The sample loaded in lane 4 was used for the experiment. PCR DNA (NEB) marker was loaded in lane 1.

**Supplemental Figure 2**

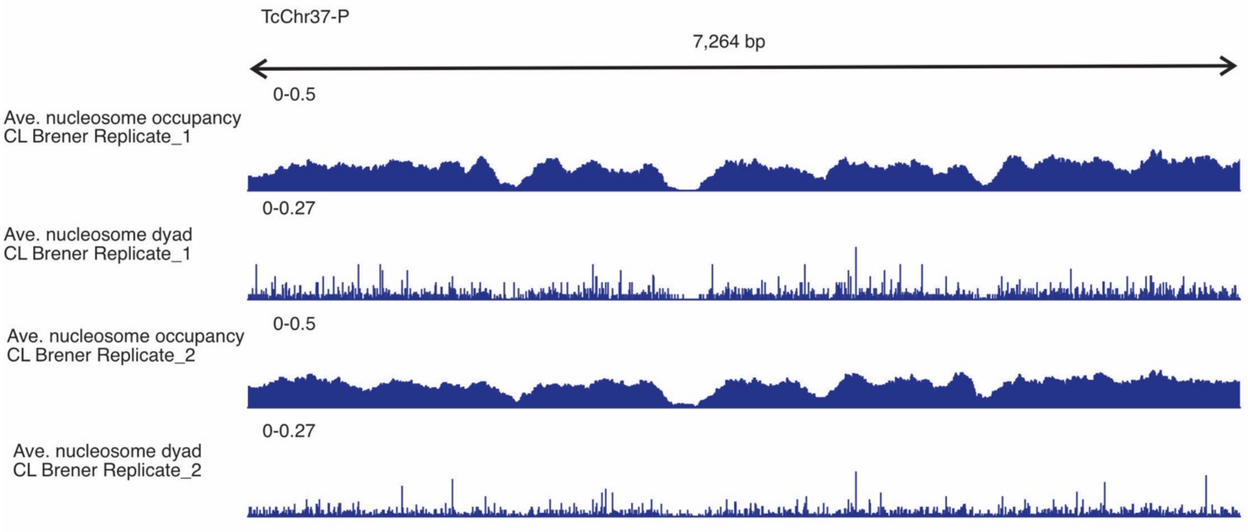


**Supplemental Figure 2. Nucleosome occupancy and nucleosome position maps**. Normalized nucleosome occupancy and nucleosome position maps are shown for a representative region of chromosome 37-P from de non Esmeraldo-like haplotype for both replicate experiments

**Supplemental Figure 3 (associated to figure 5)**

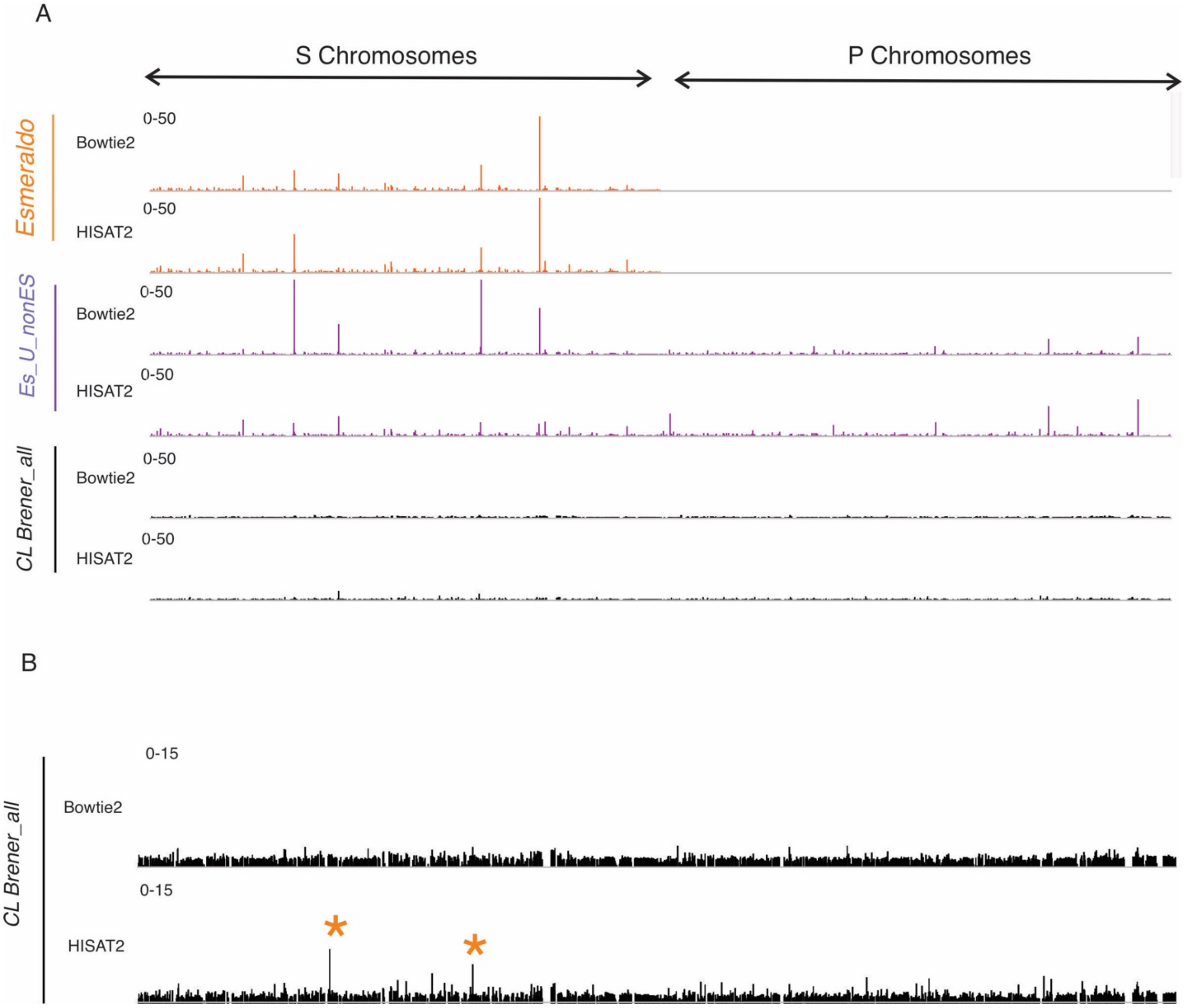


**Supplemental Figure 3. Using *CL Brener_all* genome prevents spurious alignments. (A)** IGV image for normalized nucleosome occupancy maps for the whole genome generated raw reads from one representative data set (replicate 1) aligned to the *Esmeraldo* (orange), *Es_U_nonEs* (purple) and *CL Brener_all* (black) genomes respectively either with Bowtie2 or HISAT2. **(B)** Magnified IGV image for the nucleosome occupancy maps generated from data aligned to the *CL Brener_all* genome represented in (A). Orange (*) indicates the main artefacts introduced when using HISAT2.

**Supplemental figure 4 (associated to figure 5)**

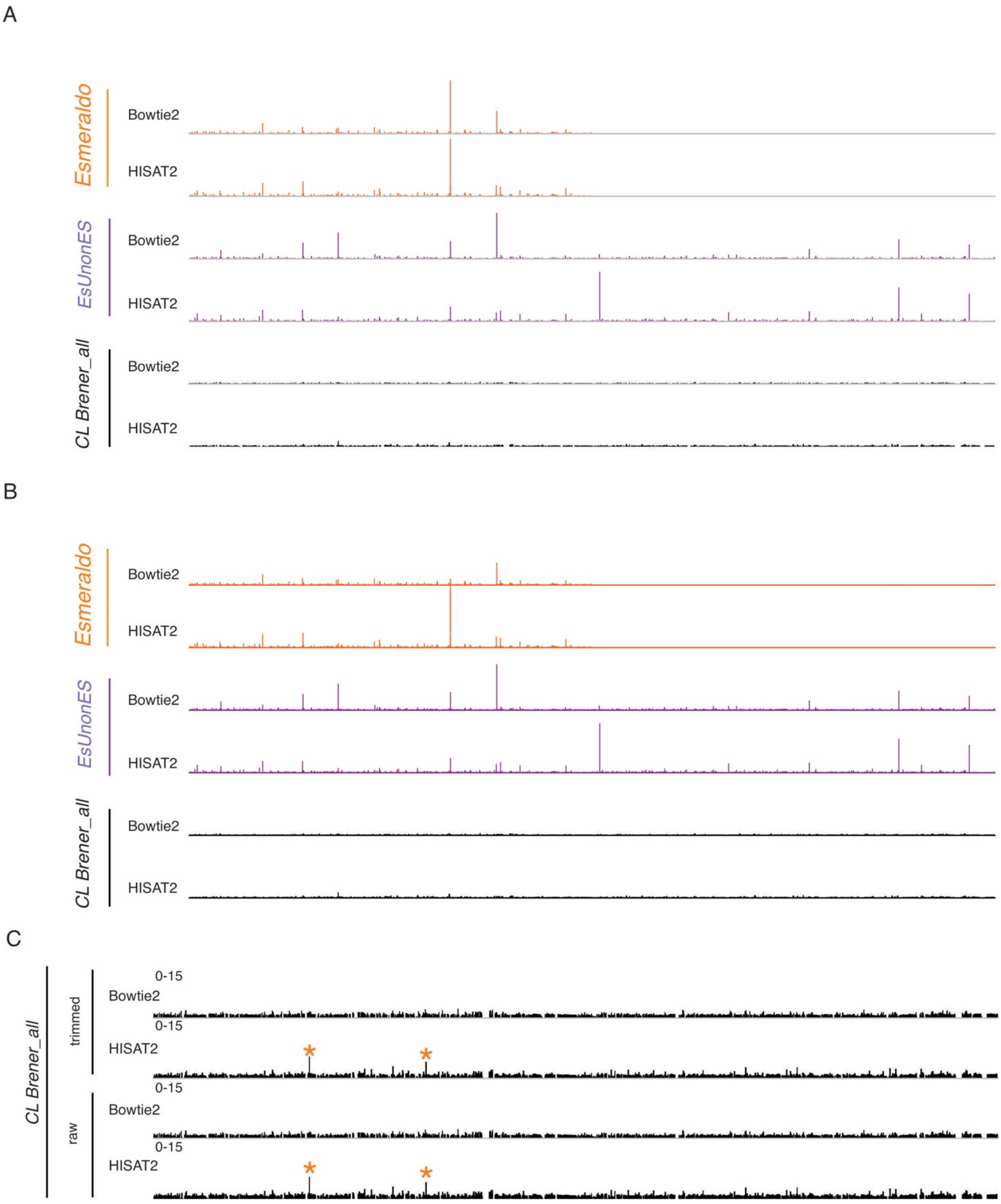

**Supplemental Figure 4. The use of *CL Brener_all* genome prevents spurious alignments.** IGV image of the whole genome for normalized nucleosome occupancy maps generated from **(A)** trimmed **(B)** and raw reads for one representative data set (replicate 2) aligned to the *Esmeraldo* (orange), *Es_U_nonEs* (purple) and *CL Brener_all* (black) genomes respectively either with Bowtie2 or HISAT2. **(C)** Magnified IGV image for the nucleosome occupancy maps generated from data aligned to the *CL Brener_all* genome represented in (A) and (B). Orange (*) indicates the main artefacts introduced when using HISAT2.

**Supplemental 5 (associated to Figure 6)**

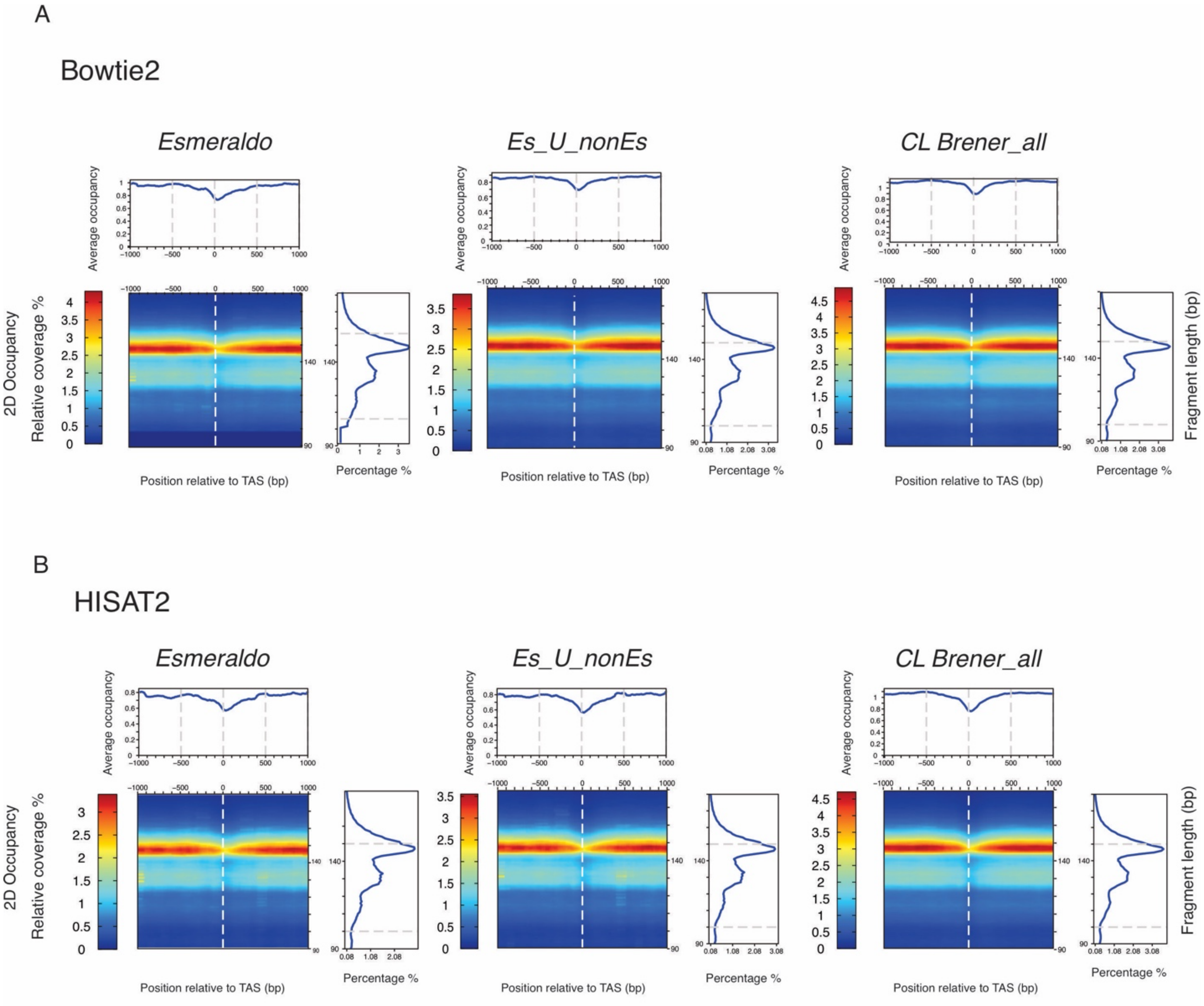


**Supplemental Figure 5. Using *CL Brener_all* genome prevents artifactual signals**. Average nucleosome occupancy and 2D occupancy plots performed from raw reads for one representative data set (replicate 1) process with either **(A)** Bowtie2 or **(B)** HISAT2. Red: High nucleosome density; blue: low nucleosome density.

**Supplemental figure 6 (associated to figure 6)**

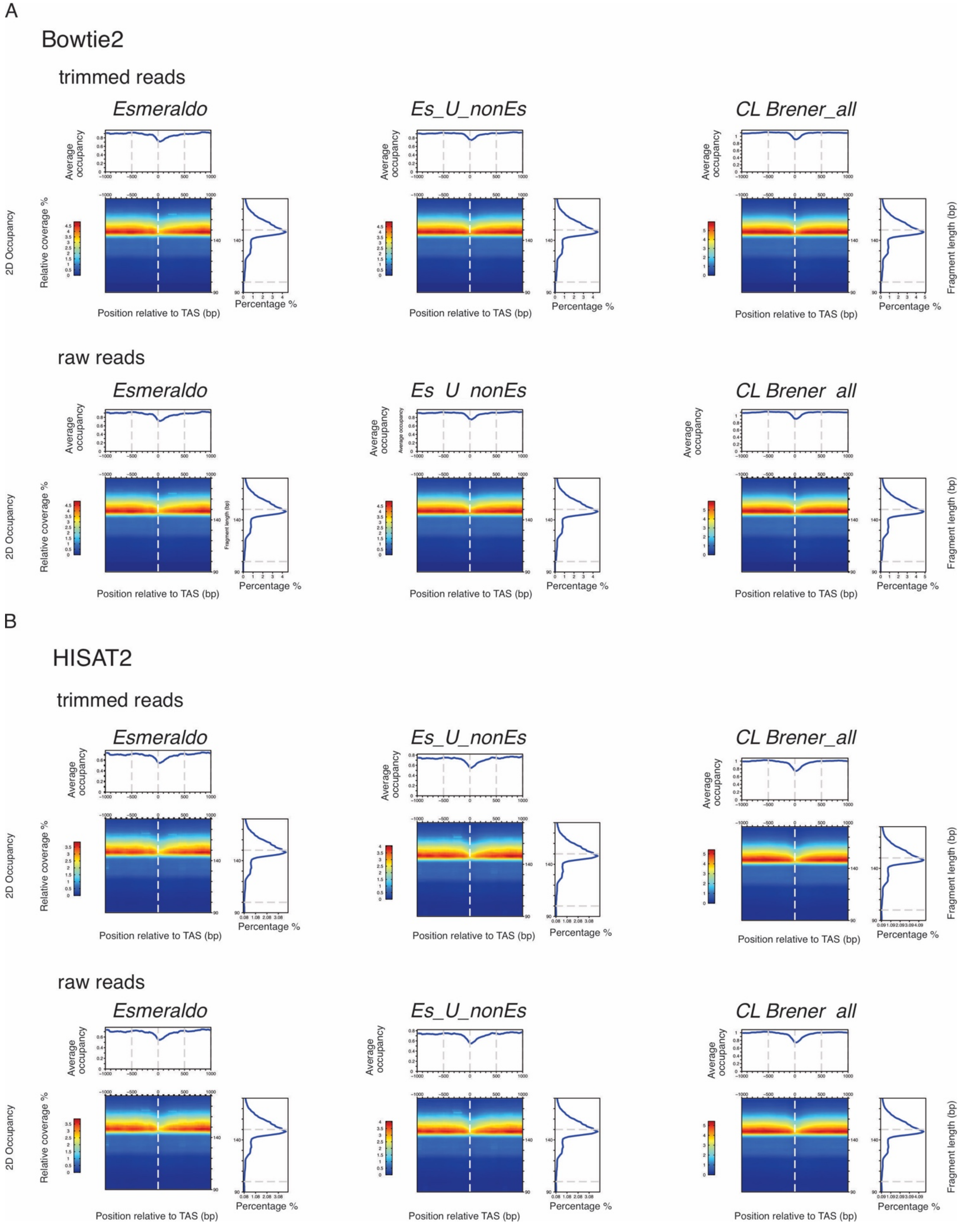

**Supplemental Figure 6. Using *CL Brener_all* genome prevents artifactual signals**. Average nucleosome occupancy and 2D occupancy plots performed from trimmed and raw reads for one representative data set (replicate 2) process with either **(A)** Bowtie2 or **(B)** HISAT2. Red: High nucleosome density; blue: low nucleosome density.

**Supplemental table 1**

| | | | | | | | |
|---|---|---|---|---|---|---|---|
| *Cl Brener* | Bowtie2 | trimmed reads | genome | 0 times | 1 time | > 1 time | overall |
| | | | *Esmeraldo* | 43 | 24.7 | 32.31 | 57 |
| | | | *EsUnonES* | 31.34 | 11.62 | 57.03 | 68.66 |
| | | | *CL Brener_all* | 23.3 | 10.3 | 66.4 | 76.7 |
| | | raw reads | genome | 0 times | 1 time | > 1 time | overall |
| | | | *Esmeraldo* | 43.98 | 24.25 | 31.76 | 56.02 |
| | | | *EsUnonES* | 32.58 | 11.41 | 56.01 | 67.42 |
| | | | *CL Brener_all* | 24.68 | 10.11 | 65.21 | 75.32 |
| | Hisat2 | trimmed reads | genome | 0 times | 1 time | > 1 time | overall |
| | | | *Esmeraldo* | 43.07 | 45.09 | 11.84 | 56.93 |
| | | | *EsUnonES* | 33.36 | 49.53 | 17.11 | 66.64 |
| | | | *CL Brener_all* | 30.96 | 44.75 | 24.28 | 69.04 |
| | | raw reads | genome | 0 times | 1 time | > 1 time | overall |
| | | | *Esmeraldo* | 44.09 | 44.29 | 11.62 | 55.91 |
| | | | *EsUnonES* | 34.55 | 48.65 | 16.8 | 65.45 |
| | | | *CL Brener_all* | 32.2 | 43.95 | 23.84 | 67.8 |

**Supplemental Table 1. Statistics of alignments.** The percentage of alignments for the different paths tested with replicate 2 is summarized. Values obtained when using trimmed reads are colored in green scale; values obtained when using raw reads are colored in yellow scale. The higher the value the darker the color.
